# Changes in ESCRT-III filament geometry drive membrane remodelling and fission *in sillico*

**DOI:** 10.1101/559898

**Authors:** Lena Harker-Kirschneck, Buzz Baum, Andela Šarić

**Affiliations:** Department of Physics & Astronomy, University College London, London, United Kingdom; Institute for the Physics of Living Systems, University College London, London, United Kingdom; MRC Laboratory for Molecular Cell Biology, University College London, London, United Kingdom

## Abstract

ESCRT-III is a membrane remodelling filament with the unique ability to cut membranes from the inside of the membrane neck. It is essential for the final stage of cell division, the formation of vesicles, the release of viruses, and membrane repair. Distinct to other cytoskeletal filaments, ESCRT-III filaments do not consume energy, but work in conjunction with another ATP-consuming complex. Despite rapid progress in describing the cell biology of ESCRT-III, we lack an understanding of the physical mechanisms behind its force production and membrane remodelling. Here we present a minimal coarse-grained model that that captures all the experimentally reported cases of ESCRT-III driven membrane sculpting, including the formation of downward and upward cones and tubules. This model suggests that a change in the geometry of membrane bound ESCRT-III filaments induces transitions between a flat spiral and a 3D helix to drive membrane deformation. We then show that such repetitive filament geometry transitions can induce the fission of cargo-containing vesicles. The mechanistic principles revealed here will enable manipulation of ESCRT-III-driven processes in cells and in guiding the engineering of synthetic membrane-sculpting systems.

## 1 Introduction

Cellular membranes require constant remodelling to allow cells to maintain homeostasis, to grow and to divide. This involves protein machines that can physically sculpt membranes in both directions, toward and away from the cytoplasm. The ESCRT-III family of proteins (endosomal sorting complexes required for transport III) is the only cellular apparatus known to deform and cut cell membranes protruding away from the cytoplasm. This is a topologically difficult transition, as the membrane needs to be deformed from the inner side of the membrane neck. ESCRT-III proteins perform this task by forming spiral/helical filaments that associate with the cytoplasmic face of biological membranes (*1–4*). This enables them to perform a wide range of membrane sculpting and snipping processes from archaeal to eukaryotic cells, such as cytokinesis (*5, 6*), multi-vesicular body formation (*7–9*), virus release (*10, 11*), membrane repair (*12*), and nuclear envelope re-sealing (*13*). Despite many attempts to use physical principles to explain how ESCRT-III performs these functions (*14–17*) it is not clear how a single protein machine has the versatility to carry out this full range of functions.

While a recent model of ESCRT-III filaments as spiral springs (*14, 18*) offers a simple way to link ESCRT-III polymer formation to membrane deformation, it is unable to explain: *i)* the sign of the deformation - spiral tension can be released in both upwards and downwards directions; *ii)* membrane scission, *iii)* the role of energy consumption via Vps4 ATPase in ESCRT-III function, and *iv)* the ability of ESCRT-III to deform both flat and curved membranes. Here, we have used coarse-grained molecular dynamics simulations to develop the first particle-based model of ESCRT-III filament function. Strikingly, our model suggests that a single filament geometry cannot explain the experimentally observed behaviours and that membrane shaping and topological transitions require energy-dependent transitions in chiral filament geometry.

## Results

### Coarse-grained Model

Our nanoscale ESCRT-III filament model is built of beads connected by springs. The minimal unit required to construct a chiral filament that preserves its flat spiral shape was found to be a triplet of beads, where the beads of neighbouring triplets are interconnected as shown in Fig. 1a. Filament geometry is controlled by bond lengths linking neighbouring triplets (Fig. 1a), while rigidity, measured by filament persistence, is controlled by bond strength between the triplets. Since building units and bond lengths between neighbouring units are equal, the target geometry of such a filament is a closed ring of radius *R*. Though some ESCRT-III filaments are observed as rings (*1,18–20*), the effects of volume-exclusion will force longer filaments to form spirals with the adoption of non-preferred curvatures causing a build-up of tension in the filament. The membrane is described using a coarse-grained one particle thick membrane model that reproduces the fluidity and correct mechanical properties of biological membranes (*21*). As a check, the main results have been also repeated using a three-beads-per-lipid membrane model that explicitly accounts for the existence of the bilayer structure (*22*) (Fig. S5). Finally, the membrane-binding face of the filament is modelled via a short-ranged attractive potential between two beads of the triplet (coloured in blue) and the membrane beads. This potential describes generic adsorption on the membrane, including screened charge-driven adsorption. Further details of the model are described in the Supplementary Information.

**Figure 1:**
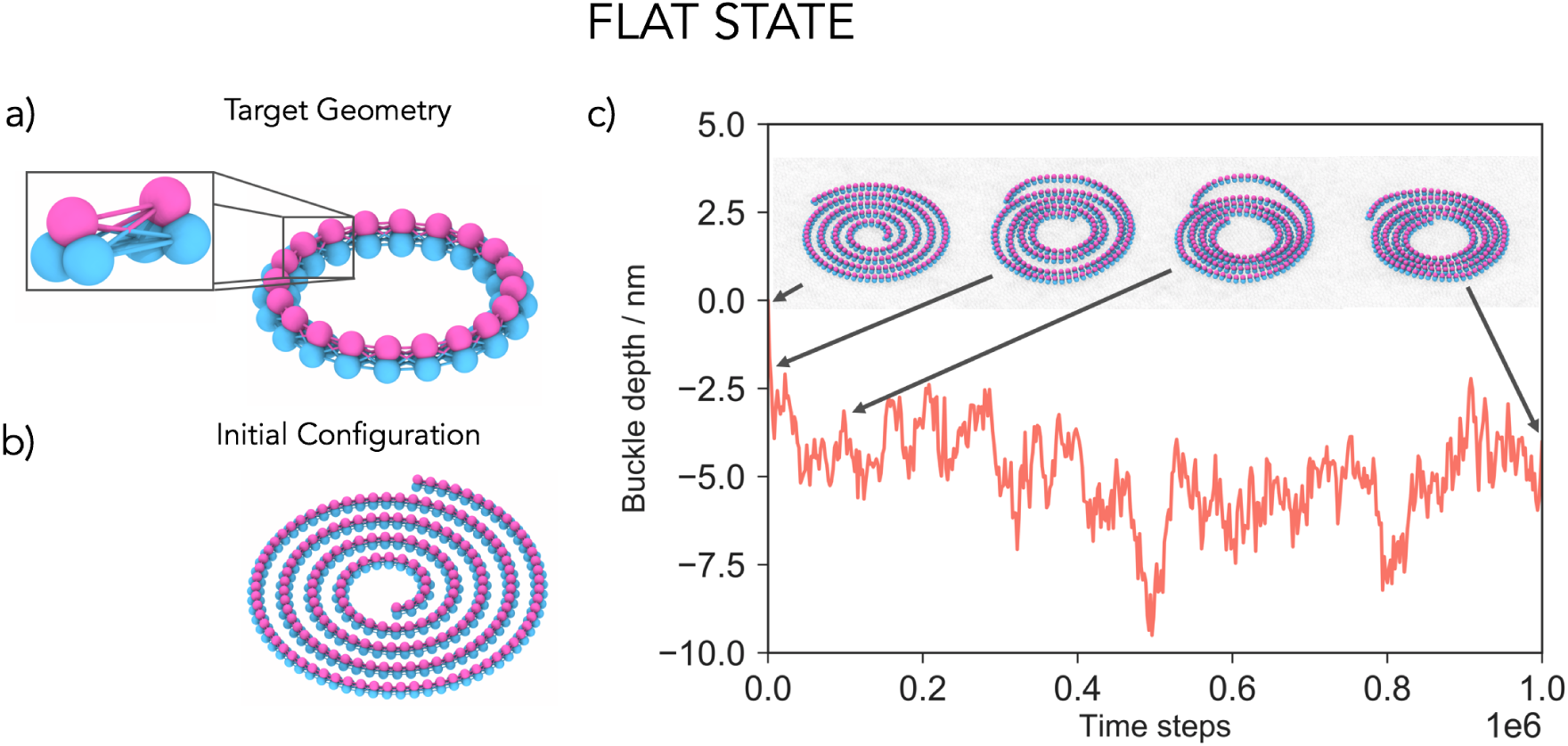
Model development. **a)** Filaments are built out of interconnected three-beaded units. The target geometry of the filament is a flat ring whose radius is determined by the rest lengths of bonds between the triplet units. The inset highlights two neighbouring units connected by 9 bonds to preserve the spiral chirality. The blue beads of the triplet are attracted to the membrane. **b)** If the filament is longer than the circumference of the target ring, it will acquire a geometry of a spiral with tense bonds, which is our initial configuration of the system. **c)** Placing a flat spiral on the membrane does not lead to a significant membrane deformation. The filament density increases, but its membrane-attracted face remains trapped in the same 2D plane. A shallow buckle that develops around −5 nm is due to the membrane enveloping the filament to maximize their contact surface. The arrows highlight simulation snapshots at specific time steps. The radius of the target geometry is *R* = 20.4 nm and the persistence length of the filament *l*_p_ = 1.8 · 10^3^ nm.

### Single filament geometry cannot deform membrane

Pre-assembled planar ESCRT-III spirals (Fig. 1b) were placed on an equilibrated flat membrane, and their behaviour followed over time as tension in the system was allowed to relax (Fig. 1c). Since the outer arms of the spiral are stretched beyond the preferred filament curvature, while the inner arms are compressed, the filament will attempt to reach mechanical equilibrium by contracting its outer and expanding its inner filament arms, as previously suggested (*14, 18*). This leads to the formation of dense spirals, since the filament cannot overlap with itself. If the modelled filament was allowed to overlap with itself, it would take on a ring shape of the target curvature (Fig. S3). Remarkably, for any tested spiral geometry or level of stored tension, spirals remained effectively flat on the membrane (Video1), even when all three beads of the filament triplets were allowed to bind to the membrane and get wrapped by it. Thus, under our model, a tense filament containing only in-membrane-plane curvature is not sufficient to drive membrane deformation. In line with this, *in-vitro* experiments have reported ESCRT-III spirals that can grow to several hundred nm in radius (*18, 23*), without deforming the membrane on their own (*1, 24*).

### Filament geometry changes

ESCRT-III filaments have also been observed in a variety of 3D shapes, such as helices and cones (*1, 16, 19, 24–28*), indicating that ESCRT-III filaments can assume alternative target geometries. To account for this, we allowed our filaments to switch between two well-defined categories of geometrical states. In the first category, the target geometry is a ring that has its membrane binding site lying in a 2D plane, leading to the formation of spirals when bound to a membrane. In the second, the target geometry switches to a ring with its membrane binding site globally rotated outwards assuming a tilt angle *τ* (Fig. 2a). The membrane attachment site is now sitting on a 3D cone/tubule surface, which lets the filament take on 3D helical shapes (Fig. 2a). We suggest that this internal filament tilt, which has not been taken into account in previous models (*14–16*), drives membrane deformation. When a planar spiral-shaped filament is placed on a flat membrane and the target geometry is switched throughout the filament into the “tilted state, we observe that the relaxation of the filament is accompanied by a clear buckle in the membrane. This deformation develops away from the cytoplasm, and grows in time to a fixed depth (Fig. 2b, Video 2).

**Figure 2:**
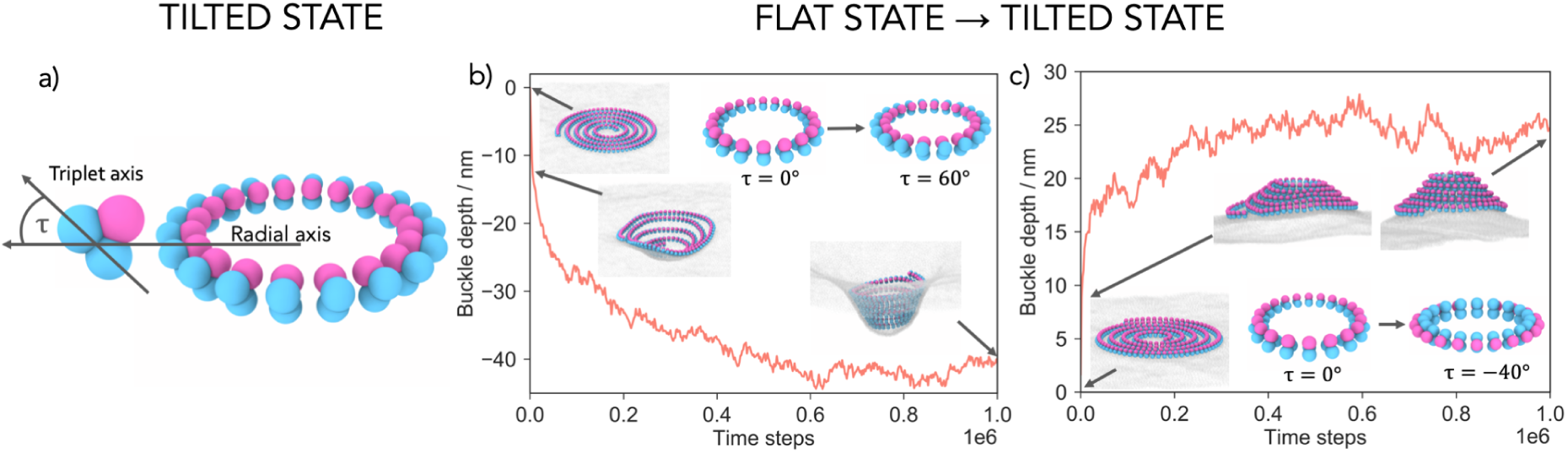
Transition between filament geometries creates membrane deformation. **a)** The tilted state is defined by an angle *τ*, which is the angle between the radial axis and the triplet subunit axis. **b)** Rotating the target geometry ring by a tilt angle *τ* (*τ* = 60°, depicted in the inset) creates downward membrane deformations, away from the cytoplasm. The curve shows the depth of the deformation over time (*R* = 15.3 nm and *l*_p_ = 1.8·10^3^ nm). **c)** The direction of the deformation can be reversed by tilting the filament outwards, rather than inwards. The new target geometry is now a ring with *τ* = −40°, *R* = 11.5 nm, and *l*_p_ = 1.8· 10^3^ nm. The underlying curve shows the height of the developing deformation over time.

This works as follows. The membrane deformation is initiated by the filament internal tilt which transforms the filament-membrane attraction site from a 2D plane to a 3D conical surface with aperture *θ* = 90° − *τ*. This initiates an out-of-plane membrane deformation, which frees the filament from being trapped in a 2D plane, allowing the filament arms to move in 3D and relax closer to their desired radius *R*. While doing so, the outer filament arms push the inner arms down into the buckle, deepening the deformation. Because this is achieved via volume exclusion, tension in the filament only makes an indirect contribution to membrane deformation by encouraging the filament to enter the buckle (see SI and Video 4 and 5 for a control experiment). The resulting filament assumes a tightly-coiled helical geometry, even though the filament does not possess a pitch in its target geometry.

While helical filaments will attempt to relax into a state in which neighbouring rings are stacked and tilted by *τ*, the wrapping of membrane about each separate ring of the spiral in the tilted state is not energetically favourable. The trade-off deformation is therefore a cone. Only for filament internal tilt of *τ* = 90° do we observe tubule formation (Fig. 3a).

**Figure 3:**
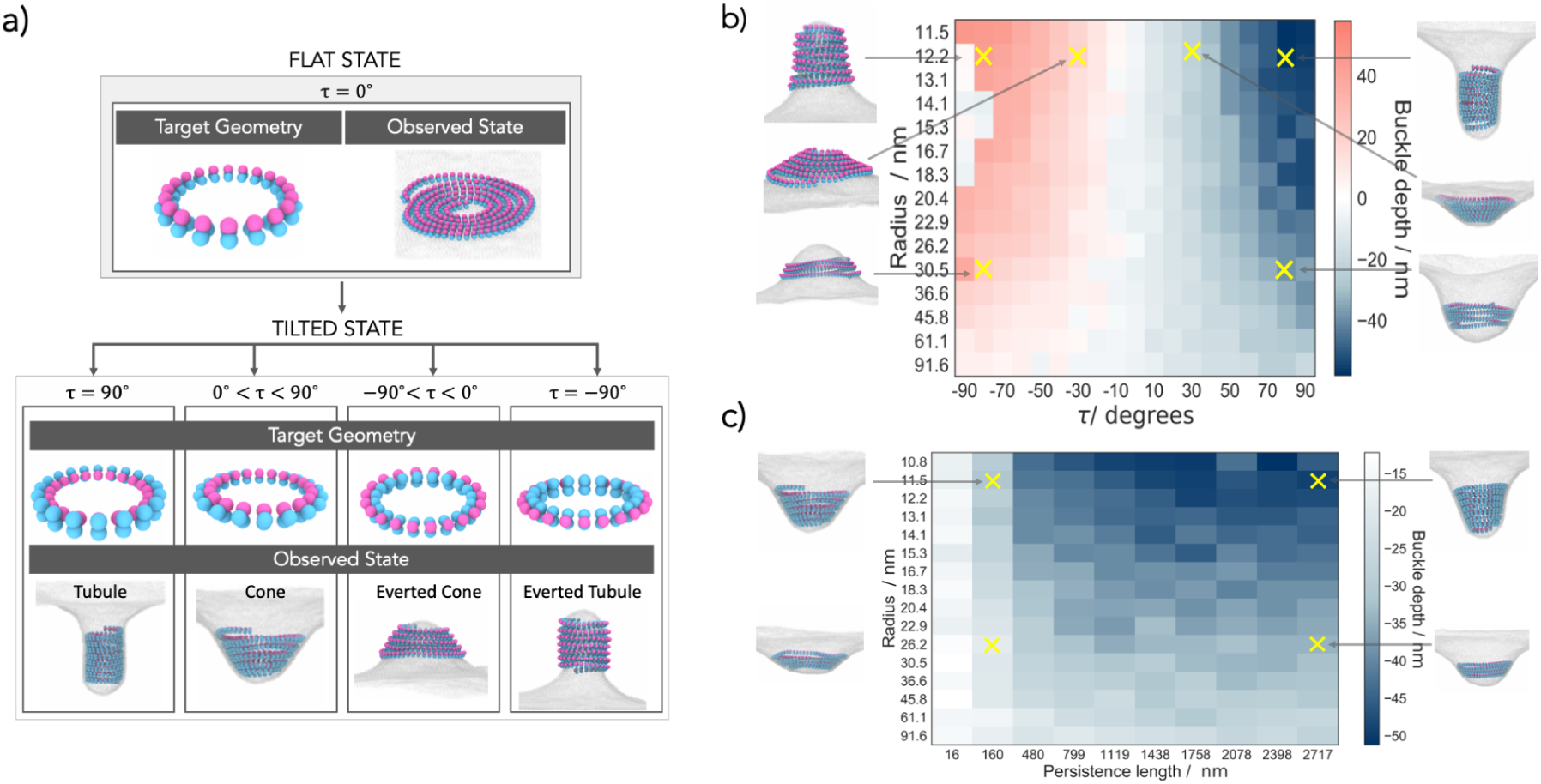
The filament tilt determines the shape and sign of the membrane deformation. Overview of the two-state model, providing examples for different outcomes. In the flat state the target geometry is a planar ring with the membrane attraction sites facing downwards (*τ* = 0°) and the filament is observed as a flat spiral. By internally rotating the target geometry ring we enter the tilted state in which tubules develop for *τ* = − 90°, cones for 0° < *τ* < 90°, everted cones for *τ* < 0°, and everted tubules for *τ* = −90°. **b)** The buckle depth dependence on the target radius *R* and the angle *τ* of the tilted state. Each simulation started off with the same initial spiral in the flat state with persistence length *l*_p_ = 1.8·10^3^ nm. **c)** The buckle depth dependence on the target radius *R* and the persistence length *l*_p_ of the tilted state (*τ* = 60°). Each simulation started off with the same initial spiral in flat state. The insets show snapshots of the representative cases.

### Quantifying membrane deformations

We found the filament tilt angle has the most impact on the shape of the deformation by determining its form and direction. This is shown in Fig. 3b, where the depth and the sign of the deformation depend on the tilt angle and the radius of the tilted state, starting from an identical initial, flat state. Strikingly, a single model parameter generates a large variety of observed states ranging from flat spirals, to conical helices of varying aperture and even tubules in both directions.

The depth of the deformation is also influenced by the preferred filament radius (Fig. 3b) and filament rigidity (Fig. 3c). Since the filaments in our simulations are of constant lengths, filaments of large preferred radii will result in shorter helices and will yield shallower deformations. The persistence length functions here as a measure of the amount of inner tension that the filament possesses. Naturally, filaments with small persistent lengths do not have a strong drive to achieve their target geometry, and cannot deform the membrane (Fig. 3c). For increasing persistence lengths, deeper buckles develop, and their shape changes from conical towards tubular. The internal filament energy now dominates over the elastic energy to bend the membrane, and a tubular deformation that accommodates the preferred filament radius is formed. Finally, to test the possible role of the helical pitch, which is not implemented in our model, we also repeated our simulations with introducing an explicit helical pitch (see SI). Interestingly, we found that it does not influence the resulting deformations and that the filament always remains in a tightly-coiled helix state when deforming the membrane (Fig. S4).

The model also provides a simple and intuitive explanation for the origin of the symmetry breaking in the membrane deformation. The buckling direction is determined by the sign of the tilt angle *τ*, i.e. by the 3D chirality in the filament. We should therefore be able to reverse the buckle direction simply by reversing the filament tilt. As can be seen in Fig. 2c and 3b by moving membrane-binding sites to the inside of the filament, buckles can be induced in the opposite direction, towards the cytoplasm (Video 3). In line with this possibility, Cullough et al. reported the formation of both upward and downward buckles of ESCRT-III filaments, depending on the filament composition (*3*). Hence it is possible that ESCRT-III filaments could induce different geometries depending on whether a spiral changes into a helix or into an inverted helix.

Interestingly, in our model there is a bias in the system that leads to a preference for downward buckles (as can be seen in Fig. 3b). Simulations for large negative tilt angles (*τ* = −80° to − 90°) do not show any deformation, an anomaly that is not mirrored for positive tilt angles. The reason for this bias in the energy landscape is that, for an upward buckle to form, the membrane must adhere to the filament from “the inside” and must adopt a larger curvature than a downward membrane deformation caused by the same amount of the filament rotation, rendering upward deformations more costly. In the case of downward deformations, the membrane envelops the filament from the outside, resulting in a smaller curvature, reducing the amount of energy required to deform the membrane. These observations may explain why cytosolic ESCRT-III filaments preferentially deform and cut membranes away from the cytoplasm.

### Membrane scission

To explore whether transitions in filament geometry can also drive scission, we introduced a simple generic cargo into the model. This cargo is allowed to weakly adhere to the membrane (see SI for details) so that it remains adsorbed and creates a shallow deformation. The adhesion is, however, too weak to cause substantial cargo wrapping by the membrane and spontaneous budding. We then polymerise a flat ESCRT-III spiral around the cargo. This models the way the ESCRT system is thought to corral cargo in cells (*29*). Under these conditions, a switch in filament geometry induced by a transition to a positively tilted state initiates a conical membrane deformation, which traps the cargo at its centre underneath the ESCRT-III helix, as shown in Fig. 4 a). In this configuration, the filament stabilises the energetically-costly membrane neck. If the filament geometry reverts back to a flat state, this destabilises the neck and causes membrane scission, releasing a membrane bud that carries the cargo particle, while the filament retracts back to the cytoplasm (Video 6). Thus, transitions between distinct geometrical states can drive ESCRT-III-mediated membrane scission to mimic the role of ESCRT-III in the formation of vesicles.

**Figure 4:**
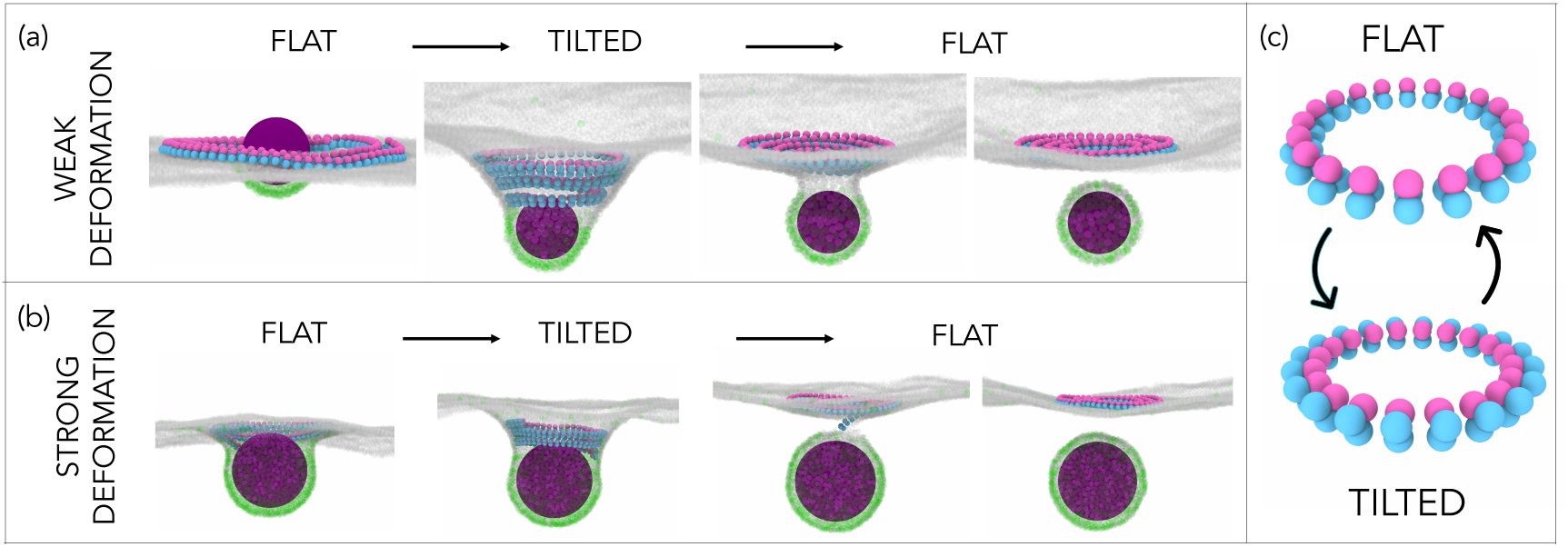
Repetitive filament transitions can sever membranes. **a)** A filament is polymerised around a cargo (magenta sphere) that weakly binds to the dark receptors in the membrane but cannot bud off on its own. Switching from a flat (*R* = 14.1 nm, *τ* = 0°) to a tilted state (*R* = 14.1 nm, *τ* = 13°) causes the membrane deformation. Switching back from the tilted to the flat state causes cargo budding where the filament is released back to the cytoplasm. **b)** The repetitive filament geometry changes also drive membrane scission in the case where the cargo has already created a substantial membrane deformation on its own, achieved by using a higher density of membrane receptors. Here the filament change from flat to helical enables the filament to enter the membrane neck and deepen it, while the opposite geometry change again performs the scission. The filament parameters and simulation protocol are the same as in (a). **c)** Target geometries of the flat and tilted states.

The mechanism by which repetitive filament geometry changes sever membranes can also be operational when substantial membrane deformation has already been formed by the cargo itself, as observed in the case Gag-driven budding of HIV-1 (*30*). Fig. 4 b) shows the scenario in which we used a larger concentration of membrane receptors such that the cargo binds to the membrane strongly enough to cause a substantial membrane deformation, but still not strongly enough to bud off on its own. In this case the helical filament geometry is needed for the filament to enter the membrane neck, while the change from helical to flat again performs the membrane scission (Video 7).

## Discussion

This minimal coarse-grained model of ESCRT-III filaments in contact with lipid membranes captures many of the experimentally observed behaviours of this versatile membrane apparatus, including different filament morphologies, diverse membrane deformations, scission, as well as cargo-loaded membrane budding and ESCRT recycling (*31*). The key ingredient of the model is the transition of the filament between two different geometrical states a flat one, where the membrane-binding sites of the filament lie on a flat plane inhibiting membrane deformation, and a helical one, where the membrane-binding sites shift into a 3D surface.

How might the different geometrical states be achieved in the context of ESCRT-III functions *in vivo*? We suggest that the presence of different members of the ESCRT-III family of proteins in copolymers (*3, 4, 25, 32*), may control the overall geometrical state of the filament. The action of Vps4 ATPase may then remodel the filament to create another geometry change, enabling cyclical geometry changes and production of mechanical work.

Our suggestion fits with the experimental evidence that binding partners of ESCRT-III can change filament structure (*3, 33–35*) and transform flat spiral filaments into helices (*1, 24*). The large variety of ESCRT-III binding partners enables the filament to move across a whole field of target geometries depending on the copolymer composition, thereby facilitating diverse ESCRT functions on very different scales and topologies. Further changes in the composition of the filament through the action of the Vps4 ATPase, which, as has been widely suggested (*24, 36*), induces filament depolymerization/turnover, would change the internal structure of the filament toward another target geometry, leading to membrane scission. We expect that similar geometrical transitions between or even within the geometry states may also enable ESCRT-III filaments to function in membrane healing and cell division. Local rather than global transitions can even lead to mixed geometry filaments that are flat on the outside and helical in the centre (*3*).

Our model makes a strong prediction that the geometry transition from spiral to helical is needed for ESCRT-III function. Indeed, ESCRT-III (co)filaments have been observed in planar forms and with helical conformations in solution and in cryo-EM (*1, 3, 25, 31*). We believe that a function role for this structural transition could be confirmed e.g. by using cryo-EM to image structures of filaments caught during or at the end of membrane deformation. While previous physical models of ESCRT-III function (*14–17*) have not included any energy input, our model suggests that the role of Vps4 ATPase likely lies in inducing a switch in the filament geometry. This analysis aligns well with the recent experiments by Goliand et al., in which the geometry change between a ring and a spiral ESCRT-III filament, caused by Vps4, is suggested to drive the constriction of the intercellular membrane bridge between two dividing cells (*27*). Similarly, Maity et al. have recently shown that Vps4 causes changes in the helical radius of ESCRT-III helical filaments *in vitro*, again suggesting that a geometry change of the filament is underlying ESCRT-III-mediated membrane remodelling (*37*). Finally, Pfitzner et al. have recently demonstrated that Vps4 ATP-ase promotes sequential changes in the composition of various ESCRT-III proteins within the filament, which is directly coupled to the filament’s ability to remodel the membrane (*28*). Hence the multiple filament geometry changes proposed by our model might be caused by an intricate interplay between the Vps4 ATP-ase and different ESCRT binding partners.

It is also important to discuss some limitations of our model. The model is coarse-grained in nature and does not capture structural details of monomers within the spiral, but only the global chiral structure of the filament as a whole. As such, a filament in our study can also represent a co-polymer made of two or more different monomer types, and we cannot make structural predictions on the sub-filament structure and inter-monomer arrangement. Our simulations were performed using pre-assembled filaments and inducing global geometry changes. This enabled us to identify the general mechanism of ESCRT-III action, however, to capture the full detailed mechanism it will be crucial to include dynamic polymerisation and depolymerisation of the filament and local changes in the geometry. This will be the topic of our future studies.

In summary, this general physical model captures a novel non-equilibrium mechanism of membrane remodelling by elastic filaments as they undergo a global change in geometry. In our simulations the energy required to drive these transitions is effectively supplied into the system by altering the filament geometry, which mimics the changes in the filament chemical composition through binding other ESCRT components or Vps4 ATPase. This coupling between the chemical composition and filament geometry produces mechanical work with which membranes can be deformed and cut. Beyond its contribution to the understanding of the ESCRT-III apparatus and membrane remodelling, our model also opens a range of possibilities for studies of membrane physics during energy-driven processes. Our results also suggest a novel way of controlling membrane remodelling in synthetic systems, applicable for instance to membrane-deformation by self-assembled DNA origami structures (*38*), where geometrical transition of the systems should be relatively straightforward to control.

## 2 Acknowledgments

We thank Jeremy Carlton, Mike Staddon, Geraint Harker, and the Wellcome Trust Consortium “Archaeal Origins of Eukaryotic Cell Organisation” for fruitful conversations. We thank Peter Wirnsberger and Tine Curk for discussions about the membrane model implementation. We acknowledge funding from BBSRC LIDo DTP (L.H.K.), the Wellcome Trust (203276/Z/16/Z) and BBSRC (BB/K009001/1) (B.B.), and the Royal Society (UF160266) (A.Š.).

## Supplementary materials

Supplementary Information.

Figures S1 to S5.

Videos V1 to V7.

References *(1-12)*.

