## Supplementary Information for "Changes in ESCRT-III filament geometry drive membrane remodelling and fission *in sillico*"

This Supplementary Information provides details on the computer model and simulation set-up (Section I), the supporting results on filament geometry when volume exclusion is turned off (Section II A and B), and the supporting results using an explicit bilayer membrane model (Section II C).

### I. Simulation details

#### A. Minimal model of ESCRT-III

We developed a minimal, coarse-grained, model of ESCRT-III based on general structural information gathered from *in-vitro* experiments [1–5]. ESCRT-III filaments form membrane-adsorbed spirals, and our simulations start with a pre-assembled, tense spiral adsorbed onto a deformable membrane. We constructed the filament model based on three distinct general properties that are observed in all ESCRT-III filaments: strong internal bonds between sub-units, an attraction site to the membrane and an intrinsic chiral curvature. It is important to note that our model does not capture the fine structural details of monomers within the spiral. It only captures the global chiral structure of the filament as a whole. We used the structure in Ref. [1] to map the simulation length-scale (size and curvature of the filament) to the physical one observed in experiments.

##### *1. Internal bonds*

Based on cryo-EM images, it has been reported that ESCRT-III (Vps32) filaments are composed of distinct densities that are separated by narrow linkers that are 3.2 nm long [1]. Hence in our model, the filament is built as a string of ESCRT-III sub-units connected to their neighbouring units by strong harmonic bonds:

$$E_{\text{bond}} = K_{\text{bond}} \cdot (r - r_0)^2 \,, \tag{S1}$$

where $K_{\text{bond}}$ is the bond strength, $r$ the distance of the two sub-units and $r_0$ the resting distance they desire to reach. This ensures that the monomers strongly attach to each other [6], which gives the filaments the ability to withstand large amounts of tension without snapping. To model filaments of different persistence lengths we used different $K_{\text{bond}}$ values.

##### *2. Chiral curvature*

ESCRT-III monomers have been found to have a preferred curvature, while also being very flexible, which enables them to adopt a large range of angles while remaining chiral [1]. However, implementing this 3D chirality into our simulations turns out to be non-trivial. We found that a minimal requirement is a triplet of beads (3D sub-units), where the beads of neighbouring triplets are all interconnected as can be seen in Figure S1.

**Single-stranded** Initial attempts to simulate this intrinsic curvature with 1D sub-units by implementing a harmonic angle potential between harmonically bonded (Equation S1) neighbouring units throughout the filament led to mixed-chirality states (see left panel in Figure S1). The angle was implemented via:

$$E_{\text{angle}} = K_{\text{angle}} \cdot (\theta - \theta_0)^2 \,, \tag{S2}$$

with $E_{\text{angle}}$ being the angle energy, $K_{\text{angle}}$ the angle strength, $\theta$ the actual angle between the two sub-units and $\theta_0$ the angle they desire to reach. However, using this potential, the next sub-unit position within the filament is not well defined, as it can be anywhere on a circle around the angle axis with that particular angle $\theta_0$ and bond distance $r_0$. To stop the filament from relaxing its tension by adopting a zig-zag pattern, we need to energetically reward one chiral angle direction, while punishing all the others. This can be achieved by implementing curvature via the filament structure itself, rather than by using a potential.

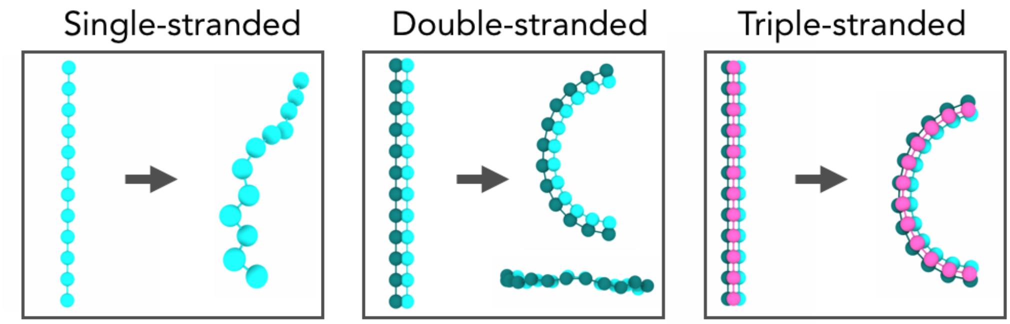


FIG. S1: Left panel: The single-stranded filament is built from one string of beads with harmonic bonds and harmonic angles implemented between consecutive beads. Since the harmonic angles can relax in multiple directions, the filament does not exhibit chirality in 3D. Middle panel: A double-stranded filament exhibits curvature when one strand has shorter bond rest lengths (here pale green) than the other (dark green). The filament in its rest state will bend in the direction of the contracting side. However, when looking at the curved filament from the side, we can see ripples forming, indicating that it is not chiral in 3D. Right panel: Adding a third stabilising string of beads on top of the other two guarantees 3D chirality given sufficient bonds are implemented between the beads.

**Double-stranded** Such an approach can be seen in the middle panel of Figure S1 with 2D sub-units. The filament is built out of two strings of beads that are rigidly attached on their sides, but have different bond rest lengths implemented along their length. We can see the resulting filament curve in the direction of the filament side that has the shorter bonds in the middle panel of Figure S1. However, looking at the system from the side reveals that it is still not chiral in 3D. The filament sub-units can move up and down in the third dimension to adjust their bond distances and relax, thereby causing un-chiral ripples (see middle panel of Figure S1).

**Triple-stranded** To achieve chirality in 3D, we need to add a third string of beads to the filament, that is sitting on top of the other two strings as can be seen in the right panel of Figure S1. It is important to realise that we are not dealing with three independent filaments, but one single filament that consists of 3-beaded sub-units (3D sub-units). The position of any bead in a sub-unit must be well-defined by the positions of the 3 beads in the previous sub-unit and their relative distances to this new bead. This means we need to define exactly 9 bonds between each set of sub-units (3 bonds per bead in a sub-unit) to specify the location of the next three beads (the next sub-unit) in 3D space. This is done by generating a sphere around each of the three original beads with the bond distance they have to a particular bead in the next sub-unit as the radius. For suitable values, these 3 spheres will cut in exactly two points in 3D space. One is already occupied by a preceding sub-unit and the other is defining the next triplet bead's position. Due to the 9 bonds, each bead from one triplet is harmonically bonded with each bead from the next triplet via Equation S1. All the bonds were given equal $K_{\text{bond}}$. For varying filament persistence length we used values $K_{\text{bond}} = 5 - 850$kT. Otherwise we used $K_{\text{bond}} = 550$ kT. If we implemented fewer than 9 bonds, the filament would lose its 3D chirality and release its energy by internally twisting and zig-zagging out of shape. Moving towards the rest length of just one bond in a forbidden way will always push the other bonds further from their equilibrium, thereby penalising that move if we sum over all bonds. We only allow the filament to deform in a chiral manner that decreases the sum of all bonds to reach its equilibrium curvature.

#### 3. *Filament structure*

The inset in Figure S2 shows the cross-section of our filament model as three beads sitting at the corners of a triangle that form a rigid object (referred to as triplet). Rigid here means, that they can't change their relative position within this triangle. What they can change is the position to their neighbouring triplets further along in the filament cross-section by adjusting their 9 interconnecting bonds.
The filament geometry is hence controlled by the bond lengths between the neighbouring triplets. The filament rigidity, measured by its persistence length, is controlled by the strength of the bonds between the triplets. Since all the sub-units are equal and hence each set of 9 bond lengths between neighbouring units are equal to the next set, the target geometry of such a filament would be a closed ring of a specific radius $R$ and angle $\alpha$ between sub-units as can be seen in Figure S2. When moving along the filament length, we keep the distance between two consecutive

centres of blue beads constant at $d_{\mathrm{const}} = 3.2\mathrm{nm}$ (equal to the measured distance between centres of densities in Ref. [1]) and define the circle radius $R$ to be the distance of the centre of the two blue beads to the centre of the circle:

$$R = ((d_{\mathrm{const}})/2)/(\sin(\alpha/2)). \tag{S3}$$

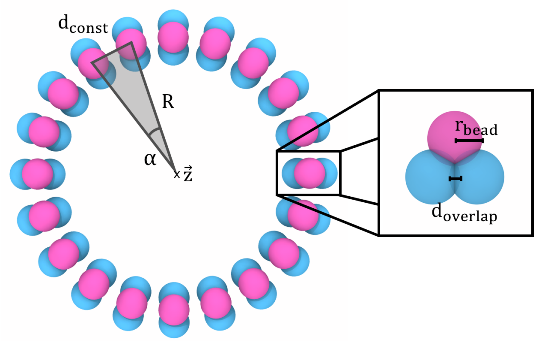


FIG. S2: The relaxed state of the filament is a ring in which all monomers reached their desired in-plane curvature, characterized by $\alpha$. The centres of blue beads have a fixed distances $d_{const} = 3.2$ nm to each other. All bead within a triplet (displayed in the inset) overlap by $r_{overlap} = 0.47$ nm. These parameters define a radius $R$ around the circle axis $\vec{z}$ via equation S3.

We simplified the filament without changing the overall behaviour by making the end of one sub-unit the beginning of the next. The three beads within the rigid object also overlap by a distance $r_{\mathrm{overlap}}$, to avoid membrane beads getting stuck between the filament beads as can be seen on the inset of Figure S2. We can calculate the bond distances of the filament's equilibrium ring state using simple geometry. We just rotate the triplet forming the beginning of the sub-unit by the angle $\alpha$ around the circle axis $\vec{z}$ at a distance $R$ to find the other end of the monomer in its relaxed state (see Figure S2). Then we can extract the 9 bond rest lengths by calculating the distances of the corresponding beads. The same applies when the filament is internally rotated by a tilt angle $\tau$ around the middle of the two blue beads. Given we define this point to mark the circle radius, it does not get affected by the tilt angle.
Unlike in the previous 1D and 2D models, here only the blue part of the filament interacts with the membrane via a Lennard-Jones potential defined in Equation S4. The pink beads are neither attracted nor repelled by the membrane. Instead they interact via volume-exclusion (a Lennard-Jones potential that is cut off at its minimum), a concept we implemented for all the particles in the simulation to stop them from overlapping with each other.
The size of the each bead within the triplet was equal to the simulation unit length $\sigma$. The beads of the triplet overlap by $r_{\mathrm{overlap}} = 0.2\sigma$ with each other, hence the width of the filament is $1.8\sigma$. It has been reported that ESCRT-III filaments are about 4.2nm thick [1]. We use this transformation to convert the simulation units of length to physical units. Hence $\sigma = 2.3\mathrm{nm}$.

#### B. Membrane model

The membrane is modelled using a coarse grained one-particle thick model [7], which we implemented in the LAMMPS Molecular Dynamics package [8]. In short, in this model each membrane bead is described by its position and an axial vector. The beads interact with a combination of an attractive potential that depends on the inter-bead distance and drives the membrane self-assembly, and an angular potential that depends on the angle between the axial vectors of neighbouring beads and mimics membrane bending rigidity. The spontaneous curvature per bead, which can be implemented via a preferred angle between two axial vectors, was kept at zero. Following the notation from the original paper [7], we chose the parameters $\epsilon_{\mathrm{bead-bead}} = 4.34k_{\mathrm{B}}T, \xi = 4, \mu = 3, r_{\mathrm{cut}}^{\mathrm{bead-bead}} = 1.12\sigma_0$, where $\sigma_0$ is the membrane bead diameter, such that our membrane is in the fluid phase [7] and its bending rigidity is $20k_{\mathrm{B}}T$ [9], which is in the regime of biological membranes. The membrane bead diameter was chosen to be equal to the simulation unit of time $\sigma$, hence it roughly corresponds to 2.3 nm .

#### C. Filament-membrane attraction

The attraction of the filament to a membrane bead is simulated by a Lennard-Jones potential acting between the two membrane-attracted beads of each triplet and a membrane particle:

$$E_{\mathrm{lj}} = 4\epsilon\left(\left(\frac{\sigma}{r_{\mathrm{lj}}}\right)^{12} - \left(\frac{\sigma}{r_{\mathrm{lj}}}\right)^{6}\right) + \beta, \tag{S4}$$

for $r_{\mathrm{lj}} \leq r_c$ and zero otherwise. $\epsilon = 3$ kT is the interaction strength, $\sigma = 2.3$ nm is the contact distance between a membrane-attracted bead of an ESCRT-III unit (represented in blue) and the interacting membrane particle, and $r_{\mathrm{lj}}$ being their actual distance. $\beta = 1$ kT is chosen so the potential becomes 0 at the interaction cutoff $r_c = 3.4$ nm. This potential is meant to represent generic surface-adsorption, including that via screened electronic charges, as suggested for ESCRT-III-membrane binding [1].

#### D. Cargo budding

To model ESCRT-III mediated budding of a weekly-adsorbing cargo (Fig. 4a in the main text) we included a generic cargo particle, described as a Lennard-Jones particle of a diameter equal to $8\sigma \approx 18$nm. The cargo particle interacts via volume-exclusion with all the other particles in the system (Section I F). In addition, we include 3% of "receptor" membrane beads that bind to the cargo via Lennard-Jones attraction described in Eq.(2) with a depth of 7 kT and are otherwise equal to the other membrane particles. After the equilibration, the simulation procedure was the following: we run the filament corralling the cargo in the flat state for $8 \cdot 10^4$ time steps, then switch the filament into the helical state for $1 \cdot 10^5$ time steps, and back to the flat state for $3 \cdot 10^5$ time steps,.

To model scission of a preformed membrane bud (Fig. 4b) we used the same procedure, albeit with a bigger cargo (a diameter equal to $16\sigma \approx 36$nm), 13% of receptor beads, and cargo-receptor binding of 3 kT. The bigger cargo creates a more stable membrane neck in which the filament can enter in a helical conformation. The system was equilibrated for $8 \cdot 10^4$ time steps, then switched into a helical state for $1 \cdot 10^5$ time steps,and then turned back into the flat state for $3 \cdot 10^5$ time steps.

#### E. Simulation set-up

We simulated a flat square portion of a membrane consisting of 7360 beads with periodic boundary conditions in a $NpH$ ensemble with pressure $p = 0$ to model a membrane with a low tension. The simulation box height was fixed at $L_z = 460$nm, and the starting dimensions of the box in the membrane plane were $\approx 184x184$nm. The ESCRT-III sub-units (triplets) are represented by rigid objects. All the particles in the system are subject to random noise implemented via a Langevin thermostat with friction coefficient set to unity: $\gamma = m/t_0$, where $m$ is a particle mass (set to unity for all the particles) and $t_0$ the simulation unit of time. The value of the simulation time-step was $0.01t_0$. All the simulations were run for $10^6$ time-steps, although equilibrium would be typically reached in $\approx 5 \cdot 10^5$ steps.

To make sure the membrane is in a relaxed state when starting our simulations, we first let it equilibrate on its own in the simulation for a million time steps. We used the LAMMPS [8] molecular dynamics simulation package [23], to intergate the equations of motion and visualize our simulation results using OVITO, an open-source analysis and visualization tool for atomistic simulations [10].

#### F. Membrane deformation simulations

For all simulations in which we studied the ability of the filament spirals to deform membranes, we placed the pre-assembled spiral model on a hexgrid membrane plane (centre of sphere distance 2.6 nm). We then let the membrane equilibrate with the filament-membrane attraction turned on. Throughout this process we did not integrate the positions of the filament. This ensured that the filament and membrane were in contact once the equlibration finished after a million time steps. We then activated the bonds between the filament sub-units which drive the filament geometry transition, and observed the system behaviour for another 1 million time steps. We measure the buckling depth by keeping track of the smallest $z$-value of any bead within the filament at any given time and comparing it to its starting position to calculate how far it moved out of position.

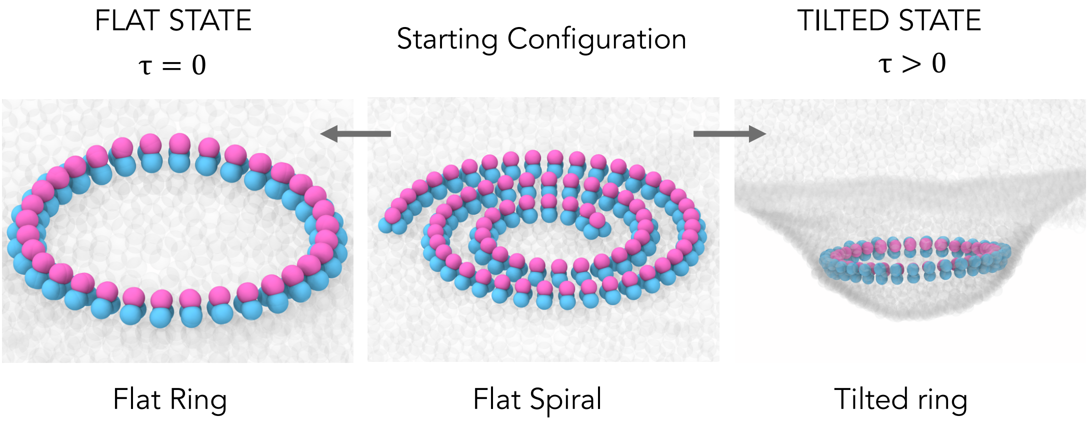


FIG. S3: Letting the filament in the middle equilibrate with volume exclusion turned off leads to the left image if the filament is in the flat state (see also Video 3) and the right image when the filament is in the tilted state (see also Video 4).

### II. Supporting Results

#### A. Turning volume exclusion off

Since the filament cannot overlap with itself the resulting geometry is a dense spiral. As a control, we repeated this simulationÂğ while allowing the filament to overlap with itself (Figure S3, Video 3-4). A self-overlapping filament in the flat state acquires a flat ring shape (left panel in Fig.S3 and Video 3). A self-overlapping filament in the tilted state acquires a tilted ring shape (right panel in Fig.S3 and Video 4).

#### B. Helical rest state

Although the filament in the "tilted state" does not possess a pitch in its target geometry, due to volume exclusion, the observed conformation will be a tightly-coiled helix. Here we test the role of imposing an explicit filament pitch in the target geometry. This would enable the filament to potentially deform the membrane even stronger.
For this purpose we have designed an experiment in which the tilted target geometry is no longer just a rotated ring, but rather a helix with a particular pitch as displayed in Figure S4 a).
In order to make our modelled filament adopt a helical shape when at rest, we introduced a small shift $dz$ along the circle axis $\vec{z}$ for each monomer (see inset in Figure S4 a). Figure S4 b) shows how the buckle depth depends on the helical rest state radius $R$ and monomer shift $dz$, which relates to the pitch $p$. Interestingly the imposed helical pitch does not influence the buckling depth, and the helical loops appear to be very close to each other in all simulations independent of the value for $dz$. This is surprising, as we would expect a helix with a large pitch to have more space between its helical loops and hence create a deeper deformation. This highlights the fact that it is the volume exclusion of filament arms within the deformation that really causes membrane deformation. The filament tension is only contributing indirectly. One other reason why we do not observe a trend for $dz$ is that the building blocks of the filament share the cost of deforming the membrane when they stay close together, and this accumulative effect outweighs the energetic benefits of releasing the pitch. When we simulate the spiral relaxing into a helical state without a membrane, we can see it release its pitch, suggesting the membrane is indeed responsible for this effect.

#### C. Explicit membrane bilayer

The membrane model we use in this work is a one-bead-per-lipid model that yields self-assembled fluid membranes of the bending rigidity that matches the one of biological membranes and can capture topological transitions such as membrane scission. The model, however, does not describe the hydrophobic and hydrophilic layers separately, and cannot capture separate local compression or expansion of one of the two sides of the membrane, which was indicated as possibly important for ESCRT-III driven membrane deformation in Ref. [2]. To capture the local effects on the

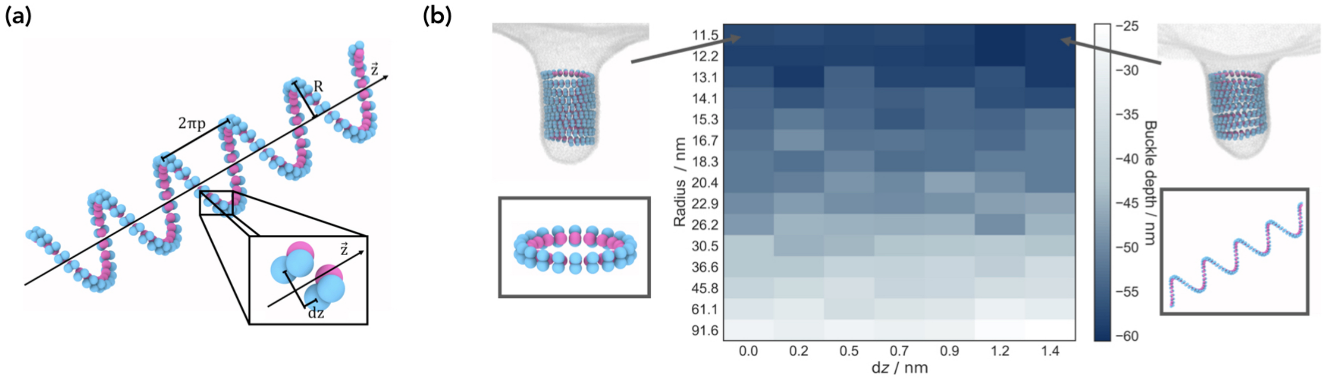


FIG. S4: **Introducing helical pitch in the tilted state**. (a) Relaxed helical filament state for radius $r = 10.3$ nm, pitch $p = 3$ nm and persistence length $l_\mathrm{p} = 1758$ nm. The zoom shows how each monomer gets shifted by $dz = 0.9$ nm along the along the $\vec{z}$ axis. The membrane-attracted blue beads are on the outside of the helix due to the positive filament tilt angle $\tau = 90°$. (b) The dependence of the depth of the deformation on the rest state radius $R$ and shift $dz$. Each simulation started off with the same flat spiral and the tilt angle is constant for all the simulations ($\tau = 90°$). Two example snapshots are displayed for different radii and $dz$ with their target geometries highlighted in rectangles beneath them.

opposite sides of the membrane bilayer, we repeated some of our key results using a coarse-grained lipid model that describes the bilayer nature explicitly [11]. In this model a lipid is described by three beads: one for the hydrophilic head and two for the hydrophobic tail. Following the original paper, we used the membrane parameters $w_c/\sigma = 1.42$ and $k_\mathrm{B}T/\varepsilon = 1.0$, which places the membrane in the fluid phase. We used 16560 lipids in the isoenthalpic-isobaric ($NPH$) ensemble with zero lateral pressure $P_x = P_y = 0$ coupled to Langevin thermostat [12]. The model for the ESCRT-III polymer remained unchanged, and the attraction between the blue bead of the spiral and the lipid head was described with the same Lennard-Jones potential as before, choosing the interaction strength of $3k_\mathrm{B}T$.

As shown in Fig. S5, when the same tense flat spiral as used in Fig. 1c was placed on the membrane bilayer the membrane deformation still failed to occur, even though the upper layer of the membrane exhibited slight local compression. The filament attraction to the bilayer remains at $\epsilon = 3$ kT. Analogously to when the one-bead-per-lipid model was used, a geometry switch from flat to tilted conformation (Fig. 2b) caused membrane deformation (Fig. S5). A repetitive geometry change (as in Fig. 4a) from flat to $\tau = 60°$ tilted and back to flat ($\tau = 0°$) around a large cargo particle with $\sigma = 22$nm still leads to membrane scission and the formation of a budded-off vesicle when performed on a bilayer membrane. The target radius of the filament spiral was kept constant at 13.1nm. In fact, all our results remained qualitatively unchanged when using the explicit bilayer model, emphasising the generality of the proposed mechanism.

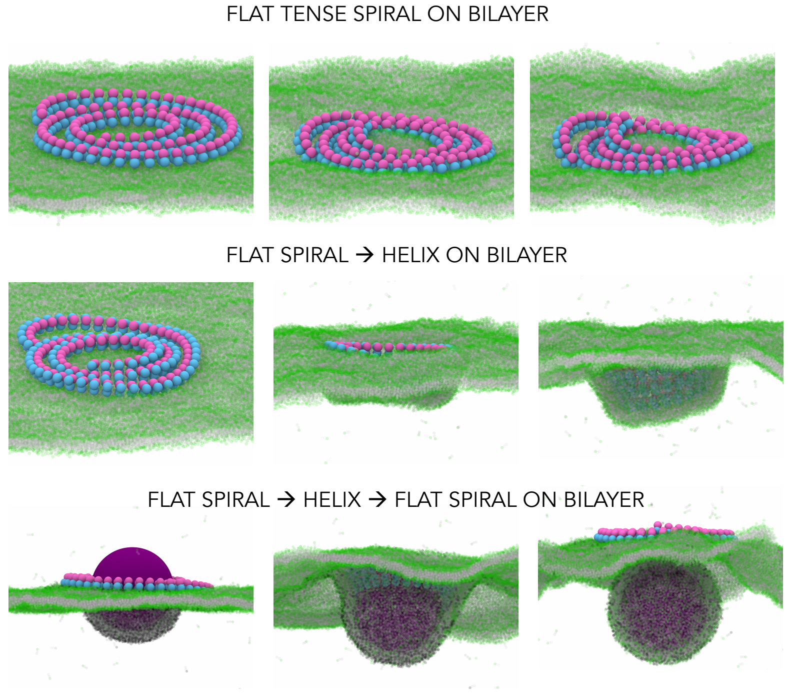


FIG. S5: **Modelling membrane bilayer explicitly using a three beads per lipid coarse-grained model**. Upper panel: flat tense spiral of the same parameters as in Fig. 1c fails to deform the bilayer. Middle panel: geometry change from flat to tilted, as in Fig. 2b, creates downward membrane deformation. Lower panel: Repetitive filament geometry changes cause membrane neck scission, as in Fig. 4a.

#### D. Supporting Videos

**Video 1:** A polymer spiral in the flat state does not lead to a significant membrane deformation. The filament density increases, but its membrane-attracted face remains trapped in the same 2D plane (see also Fig. 1c).
**Video 2:** Switching from the flat to a ring rotated by a tilt angle $\tau$ ($\tau = 60°$) creates downward membrane deformations, away from the cytoplasm (see also Fig. 2b).
**Video 3:** Self-overlapping filament spiral in the flat state ends up in a flat ring conformation (see also Fig. S3).
**Video 4:** Self-overlapping filament spiral in the tilted state ends up in a tilted ring conformation (see also Fig. S3).
**Video 5:** Switching from the flat to tilted ring ($\tau = -40°$) creates upward membrane deformations, toward from the cytoplasm (see also Fig. 2c).
**Video 6:** A cargo particle is corralled on the membrane by the spiral ESCRT-III polymer in the flat state. The2 cargo binds to the dark gray receptors in the membrane but cannot bud off on its own. Switching the filament geometry from a flat ($R = 12.2$ nm, $\tau = 0°$) to a helical state ($R = 12.2$ nm, $\tau = 60°$) deforms the membrane, where the filament stabilises the neck. Switching back from the helical to the flat state causes cargo budding where the filament is released back to the cytoplasm (see also Fig. 4a).
**Video 7:** A cargo particle creates deep membrane deformation on its own, due to the higher perfentage of membrane receptors than in Video 6. Switching the filament geometry from a flat ($\tau = 0°$) to a tilted state ($\tau = 60°$) enables the filament to enter the preformed membrane neck, deepening and stabilising it. Switching back to the flat filament state causes cargo budding where the filament is released back to the cytoplasm (see also Fig. 4b).
